# Entropy-based integration index for quantifying network integration in resting-state functional MRI

**DOI:** 10.64898/2026.06.22.733307

**Authors:** Poulami Kar, Dipayan Roy, Bhoomika R. Kar

**Author notes:** Correspondence: Poulami Kar, Centre of Behavioural and Cognitive Sciences (CBCS), Senate Campus, University of Allahabad, Prayagraj, Uttar Pradesh, India.

## Abstract

Independent component analysis (ICA) is widely used in resting-state fMRI to identify large-scale functional networks; however, existing approaches provide limited means of quantifying how network representations are distributed across independent components. We introduce an entropy-based network integration framework that characterizes the organizational architecture of canonical resting-state networks by quantifying the distribution of ICA-derived functional contributions within Yeo atlas networks. Spatial overlap between independent components and network templates is normalized to generate a probability distribution, from which Shannon entropy and a normalized integration index are derived. The resulting metric provides a continuous measure of network representational integration, ranging from specialized configurations dominated by a small number of components to distributed configurations involving multiple functional modes.

The framework was evaluated and validated using resting-state fMRI data from healthy controls, Parkinson’s disease patients with normal cognition, and Parkinson’s disease patients with mild cognitive impairment. Global entropy and integration measures were complemented by network-specific analyses, dominance profiling, principal component analysis (PCA), and multivariate centroid-distance assessments. The proposed framework revealed selective alterations in Ventral Attention and Limbic network organization associated with cognitive-status differences, while preserving overall within-group heterogeneity. Group-wise PCA independently further identified these networks as major contributors to altered network organization, and centroid-distance analyses demonstrated that observed differences reflected coherent shifts in network architecture rather than increased variability.

By quantifying the distribution of network representations across ICA-derived functional modes, this framework provides a simple, interpretable, and generalizable measure of large-scale brain organization, offering a complementary approach for studying network reorganization in health and disease.

## 1. Introduction

Resting-state functional magnetic resonance imaging (rs-fMRI) is widely to study large-scale brain organization, and Independent Component Analysis (ICA) is valuable for identifying spatially independent functional networks without predefined regions of interest. However, ICA-based analyses largely remain descriptive, focusing on component identification and spatial overlap, and provide limited means of quantifying how within and between network representations are distributed across functional components within individuals.

Current approaches for characterizing large-scale brain organization primarily focus on connectivity strength, network topology, temporal dynamics, or directed interactions. Functional connectivity analyses quantify statistical relationships between brain regions, graph-theoretical approaches characterize topological properties such as efficiency and modularity, dynamic connectivity methods examine fluctuations in network interactions over time, and effective connectivity approaches estimate the directionality or causal influence of interactions among brain regions or networks. But they do not directly quantify how canonical networks are represented across the spatially independent components identified through ICA, also referred to as functional modes. Consequently, an important aspect of network organization remains relatively unexplored: how broadly or narrowly a network is represented within the functional decomposition of the brain.

Quantifying this specialized versus distributed representation may enrich studies on healthy brain organization for learning and development - for example, longitudinal studies could test whether practice, maturation, or changing cognitive demands are associated with increased specialization of particular networks, broader distributed integration, or shifts in representational balance across functional modes - while also offering insight into clinical network organizations. In Parkinson’s disease (PD), for example, disruption of basal ganglia–thalamocortical circuitry has been proposed to increase reliance on alternate cerebellar–thalamocortical pathways in contexts involving motor timing, rhythmic cueing, and executive control (Lesiuk et al., 2018). Similarly, spontaneous or rehabilitation-related recovery in stroke may involve reorganization of large-scale functional networks, particularly motor and interhemispheric connectivity (Park et al., 2011). These examples illustrate that clinically relevant reorganization may involve not only altered connectivity strength or timing, but also redistribution of functional representations across large-scale systems. Thus, measuring how concentrated or distributed network representations are may provide complementary insight into adaptation, compensation, and disease-related reconfiguration.

To address this gap, we introduce an entropy-based framework that quantifies how canonical Yeo networks are represented across ICA-derived functional components (Yeo et al., 2011). For each network, spatial overlap between ICA components and Yeo templates is transformed into a probability distribution describing the relative contribution of individual components. To quantify the diversity of this distribution, Shannon entropy is used to derive a normalized integration index. Conceptually, low values indicate representation by a few dominant components, whereas high values indicate broader distribution and reflects greater representational integration. Unlike existing entropy-based neuroimaging measures, which typically focus on regional BOLD dynamics or functional activity fluctuations (McIntosh et al., 2010; Garrett et al., 2011; Wang et al., 2014; Nomi et al., 2016; Niu et al., 2018), the proposed framework applies entropy to ICA-derived network representation to quantify how canonical networks are distributed across functional modes, offering a complementary perspective of network representational architecture beyond connectivity strength, graph topology, or signal complexity (Bullmore & Sporns, 2009; Bassett & Sporns, 2017). Thus, it may help distinguish network reconfiguration that reflects increased specialization, broader integration, or altered representational balance, even when conventional connectivity measures identify only changes in coupling or temporal fluctuation.

Application to healthy controls (HC), PD with normal cognition (PD-NC), and PD with mild cognitive impairment (PD-MCI) patient groups showed that the proposed metric detects coherent group-level differences in network organization, particularly within Ventral Attention and Limbic systems, while remaining robust to changes in overall network heterogeneity. This suggests that the framework may provide a simple and interpretable tool for studying large-scale functional network organization across healthy and pathological states.

## 2. Methods

### 2.1 Mathematical background of entropy-based integration index

#### 2.1.1 ICA–Network Overlap

For each independent component *k*, spatial overlap with each of the seven canonical networks was quantified as the mean Z-statistic within each network mask:

*_k_*_,*i*_ where *i* ∈ {1,…, 7} indexes the seven canonical Yeo networks.

#### 2.1.2 Network Dominance

A dominant network was assigned to each independent component based on network with the maximum spatial overlap:

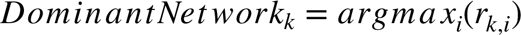

where *Dominant Net work_k_* denotes the canonical network assigned to independent component *k*. This provided a categorical representation of network specificity.

#### 2.1.3 Entropy-Based Integration Index

To capture distributed contributions across networks, the ICA-network overlap values were converted into a probability distribution:

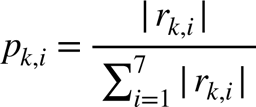

where *p_k,i_* denotes the normalized contribution of Yeo network *i* to independent component *k*, and *r_k,i_* represents the spatial overlap between independent component *k* and Yeo network *i*. Absolute values were used to ensure that both positive and negative contributions reflect the magnitude of overlap, thereby avoiding cancellation effects during normalization.

Shannon entropy (1948) is then computed as:

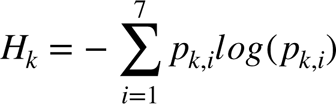

where *H_k_* denotes the entropy of independent component *k*, and the summation is performed across the seven canonical Yeo networks. The logarithm was taken as the natural logarithm.

To ensure comparability across components, entropy was normalized by its maximum possible value:

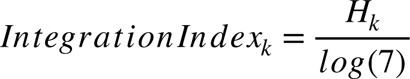

where *Integration Index_k_* denotes the normalized integration index for independent component *k*, *H_k_* is the entropy value for that component, and *log*(7) represents the maximum possible entropy when overlap is uniformly distributed across the seven Yeo networks. This yields a bounded measure between 0 and 1, where values approaching 0 indicate that the component is dominated by a single canonical network, reflecting greater specialization, and values approaching 1 indicate that the component has broadly distributed overlap across multiple networks, reflecting greater integration.

#### 2.1.4 Network-Specific Entropy and Integration Analysis

Following computation of the global entropy-derived integration index, network-specific integration indices were additionally estimated for each of the seven canonical Yeo networks to characterize the extent to which individual networks were distributed across the full set of ICs identified for a participant.

For a given network, overlap values across all ICs were normalized to form a probability distribution:

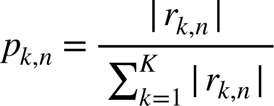

where *p_k,n_* denotes the normalized contribution of IC *k* to Yeo network *n*, *_k,n_* represents the overlap value between IC *k* and network, denotes the total number of ICs for a participant, and the denominator represents the summed absolute overlap of network across all components. Shannon entropy for each network was then computed as:

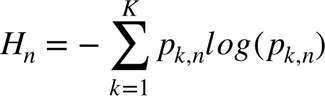

where *H_n_* denotes the entropy of network *n*, and the summation is performed across all components for that participant.

Entropy values were normalized by the maximum possible entropy for the corresponding number of ICs:

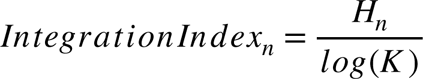

where *Integration Index_n_* denotes the normalized network-specific integration index for network *n*, and *log*(*K*) represents the maximum possible entropy when network is uniformly distributed across all *K* components.

This yielded a bounded network-specific integration index ranging from 0 to 1. Higher values indicated that a network exhibited more distributed overlap across multiple ICs, whereas lower values reflected relatively focal or specialized organizational structure.

Network-specific entropy measures were included to complement the global integration index by characterizing the organizational distribution of individual canonical networks across the ICA decomposition. Whereas the global integration index quantifies the overall degree of distributed versus specialized organization of a component, network-specific integration indices show whether particular systems (e.g., somatomotor, frontoparietal, limbic, or default mode networks) are represented relatively focally or broadly across ICs, thereby providing additional insight into the balance within/between network specialization and large-scale integrative organization.

The conceptual distinction between specialized and distributed network representations, together with the full ICA–Yeo overlap, entropy normalization, and network-specific integration workflow, is illustrated in Figure 1.

**Figure 1.**
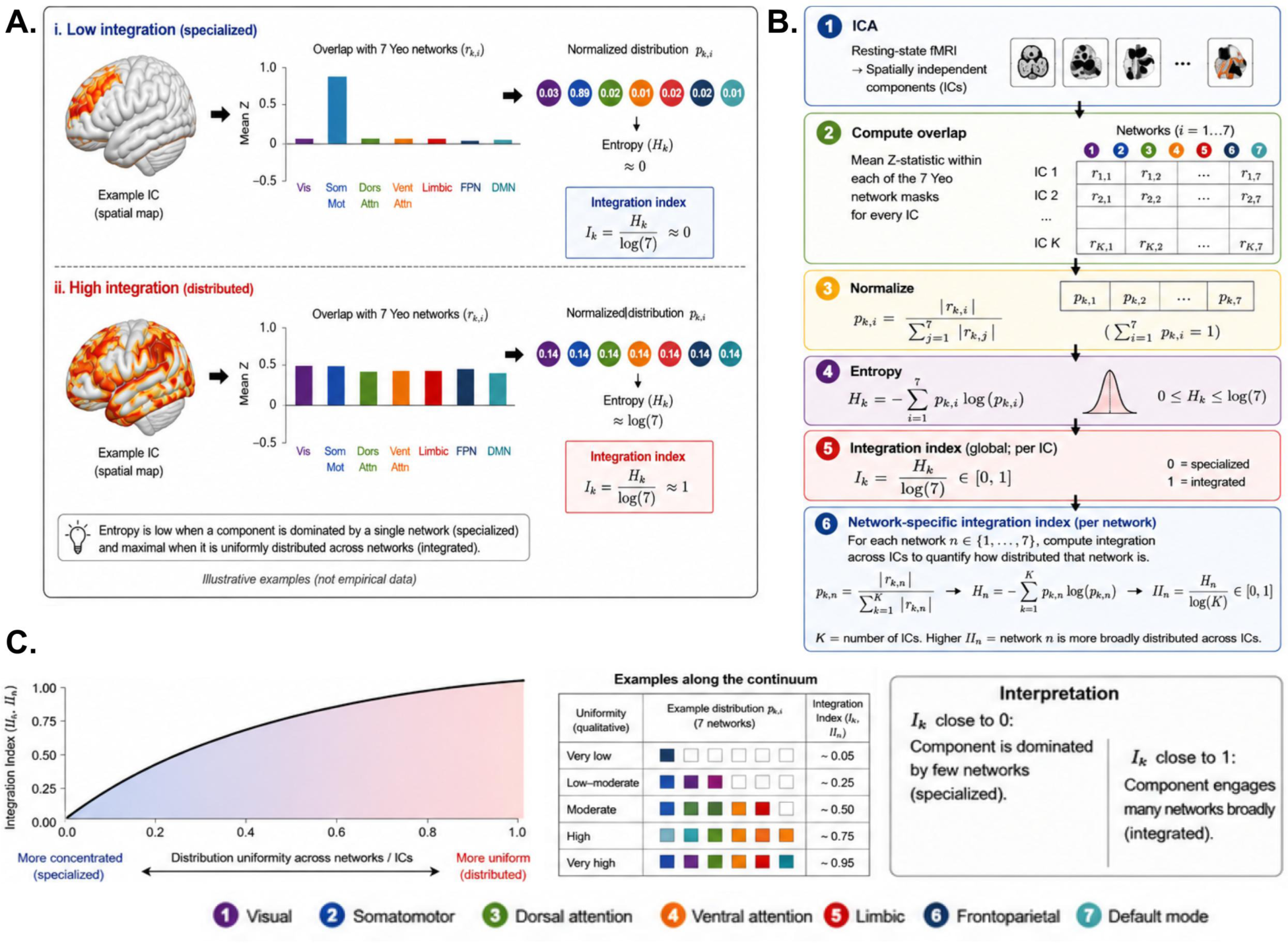
Conceptual illustration and computational workflow of entropy-based network integration index. (A) Comparison of low and high integration profiles based on the distribution of ICA–network overlap, illustrating specialization versus distributed organization. Components with overlap concentrated within a single network exhibit low entropy and integration values approaching zero, reflecting strong network specialization. In contrast, components with more uniform overlap across networks exhibit higher entropy and integration values approaching one, reflecting distributed functional organization. (B) Schematic pipeline depicting ICA decomposition, overlap computation with Yeo networks, normalization to probability distributions, and entropy-based integration index calculation. (C) Conceptual relationship between distribution uniformity and integration index, demonstrating a monotonic increase in integration with increasing uniformity across networks.

### 2.2 Application to OpenNeuro Parkinson’s Disease Dataset

To evaluate the proposed entropy-based integration index in a clinical context, the framework was applied to an openly available OpenNeuro rs-fMRI dataset (accession: ds005892) including HC, PD-NC, and PD-MCI participants (Kemp et al., 2025). This application served as an initial group-level evaluation of the proposed measure while maintaining methodological compatibility with standard ICA-based rs-fMRI workflows.

Preprocessed rs-fMRI data underwent ICA using using FSL MELODIC with automatic dimensionality estimation, allowing the number of independent components to be estimated from the intrinsic structure of each participant’s rs-fMRI data rather than imposing a fixed model order. The resulting ICA-derived spatial components were then used for subsequent overlap analysis with the Yeo seven-network atlas, and normalized entropy and the integration indices were computed for each participant.

Based on prior evidence of disrupted large-scale functional integration in PD-MCI (Baggio et al., 2015), group comparisons were used to assess whether the metric could detect altered network integration profiles in PD-MCI relative to PD-NC and HC, providing an initial evaluation of its sensitivity to PD-related cognitive-status differences. This preliminary validation framework is intended to assess the translational applicability of the proposed measure while maintaining methodological simplicity and compatibility with standard ICA-based rs-fMRI workflows.

### 2.3 Statistical Analysis

All statistical analyses were performed on network-specific integration indices derived from the entropy-based ICA–Yeo mapping framework. Statistical calculations and visualizations were carried out in RStudio using R. Analyses were designed to characterize both between-group differences in network organization and within-group patterns of network integration across HC, PD-NC, and PD-MCI. Group differences in global entropy, global integration index, and network-specific integration indices were initially assessed using one-way ANOVA. Where significant group effects were identified, pairwise post-hoc comparisons were conducted to determine the specific group contrasts driving the observed effects. This approach allowed network-level differences across HC, PD-NC, and PD-MCI groups to be examined, with particular emphasis on PD-NC versus PD-MCI comparisons to assess cognitive-status-related differences within PD.

To determine whether group differences in network integration were driven by increased heterogeneity, within-group variability was assessed using coefficients of variation (CV) for network-specific integration indices. Because integration indices can differ in their absolute magnitudes across networks, the CV provided a normalized measure of dispersion that enabled comparison of variability across networks and groups. Radar plots were used to visualize the relative inter-individual variability of the seven Yeo networks within each diagnostic group.

Group-wise Principal Component Analysis (PCA) was performed separately within each group using the seven network-specific integration indices to explore dominant patterns of network organization. The first principal component (PC1) was extracted for each group, and network loadings were examined to identify the networks contributing most strongly to the dominant organizational axis. Separate PCA analyses within each group enabled the assessment of whether cognitive-status differences were associated with a reconfiguration of the underlying architecture of network integration rather than merely changes in individual network values. Bar plots were used to visualize differences in network contributions across groups.

Multivariate heterogeneity was assessed using distance-to-centroid analysis. For each participant, Euclidean distance was calculated between their seven-network integration profile and their group centroid, providing a measure of overall deviation from the typical group profile. Unlike network-specific variability measures, centroid distance captures overall profile dispersion across all networks simultaneously, and therefore serves as an indicator of multivariate heterogeneity. Group differences in centroid distance were tested non-parametrically to determine whether integration differences reflected genuine shifts in organizational architecture or simply increased group-variability.

Together, these approaches provided a complementary and multidimensional assessment of large-scale functional network reorganization across HC and clinically-defined cognitive-status groups in PD.

## 3. Results

### 3.1 Pairwise Group Comparisons of Network-Specific Integration

Selective group differences in network-specific integration was observed. Significant effects were observed for the Default Mode Network (DMN), Frontoparietal Network (FPN), Ventral Attention Network (VAN), and Limbic Network (LIM), whereas the Visual, Somatomotor, and Dorsal Attention networks (DAN) did not show significant group-differences (Figure 2).

**Figure 2.**
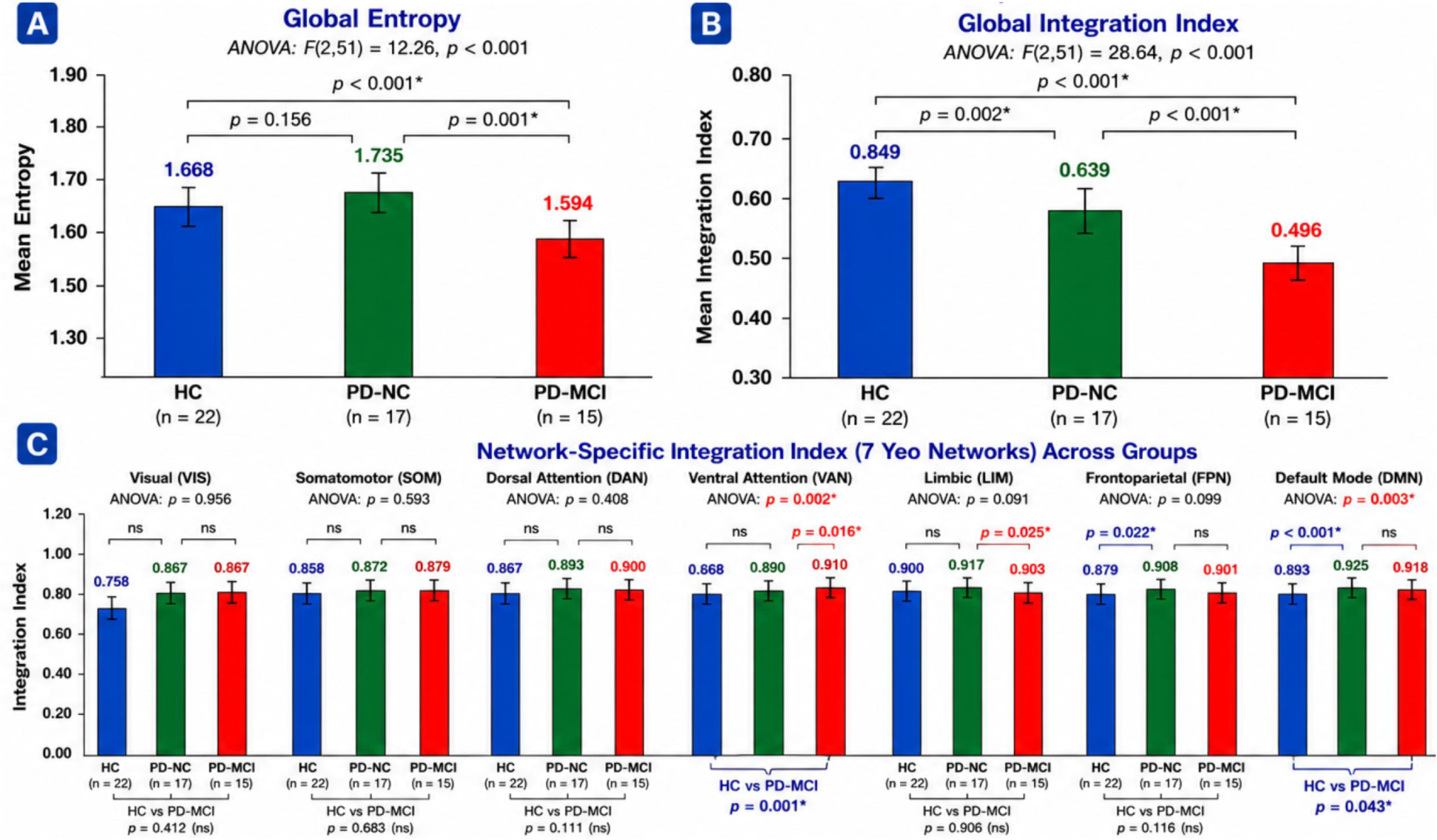
Global and network-specific entropy-derived integration indices across groups. (A) Global entropy differed significantly across HC, PD-NC, and PD-MCI groups. Post-hoc comparisons showed significantly lower global entropy in PD-MCI relative to both HC and PD-NC, whereas HC and PD-NC did not differ significantly. (B) Global integration index also differed significantly across groups, showing a graded reduction from HC to PD-NC to PD-MCI. (C) Network-specific integration indices across the seven Yeo networks revealed selective group differences. Significant omnibus effects were observed for the VAN and DMN, with trend-level effects in the LIM and FPN. Post-hoc comparisons indicated higher VAN integration in PD-MCI relative to HC and PD-NC, lower LIM integration in PD-MCI relative to PD-NC, higher FPN integration in PD-NC relative to HC, and higher DMN integration in both PD-NC and PD-MCI relative to HC. Pairwise tests were adjusted using Bonferroni correction. Bars indicate mean±SD.

Compared with HC, PD-NC participants showed greater FPN (*p*=0.022) and DMN (*p*<0.001) integration, while PD-MCI participants showed greater DMN integration (*p*=0.043). No significant HC versus PD-NC or HC versus PD-MCI differences were observed in the Visual, Somatomotor, DAN, VAN, or LIM networks.

Direct comparison of the two PD groups revealed higher VAN integration (*p*=0.016) and lower LIM integration (*p*=0.025) in PD-MCI. No significant differences were observed in the Visual, Somatomotor, DAN, FPN, or DMN networks.

Hence, group differences in network-specific integration were not distributed uniformly across all canonical networks. Instead, differences relative to HC were primarily observed in DMN and FPN integration, whereas differences between PD-NC and PD-MCI were localized to VAN and LIM integration.

### 3.2 Group-Wise Principal Component Analysis

PC1 loadings revealed distinct patterns of network organization across groups (Table 1; Figure 3). In HC, PC1 was characterized by strong loadings from the VAN (loading=−0.504), FPN (loading=−0.501), DMN (loading=−0.433), Somatomotor (loading=−0.356), and Visual (loading=−0.334) networks. The LIM showed a modest loading (loading=0.168), indicating a comparatively weaker contribution to PC1.

**Figure 3.**
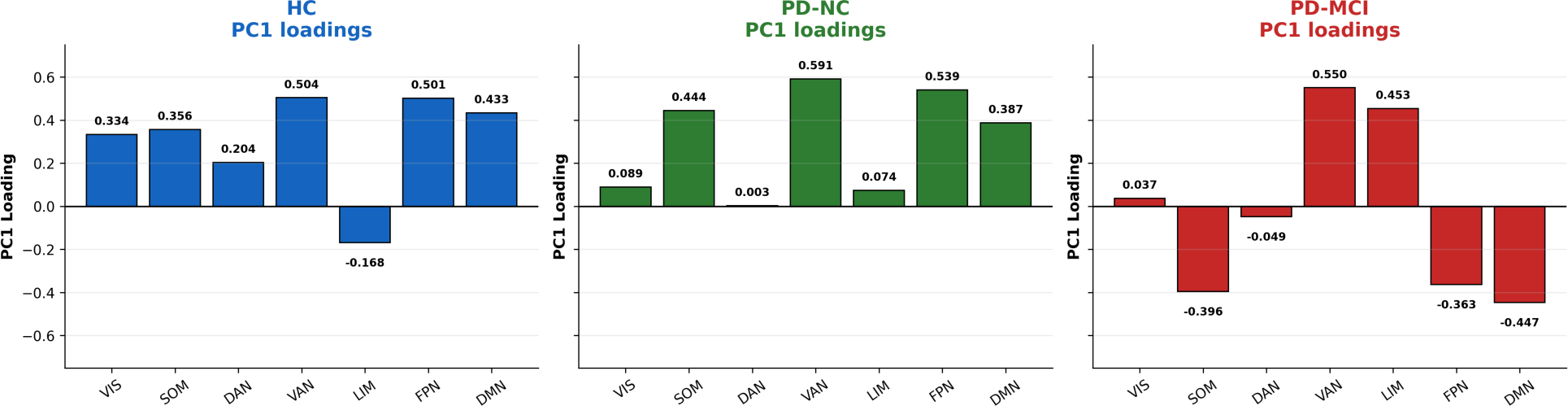
Group-wise principal component analysis of network-specific integration profiles. PC1 loadings from separate PCA analyses performed within HC, PD-NC, and PD-MCI groups using the seven Yeo network-specific integration indices as input variables. In HC, the largest absolute loadings were observed for VAN, FPN, DMN, SOM, and VIS, with a smaller contribution from LIM. In PD-NC, PC1 was primarily characterized by VAN, FPN, SOM, and DMN contributions, while VIS, LIM, and DAN showed smaller loadings. In PD-MCI, VAN and LIM emerged as the strongest positive contributors to PC1, with additional negative contributions from DMN, SOM, and FPN.

**Table 1.**
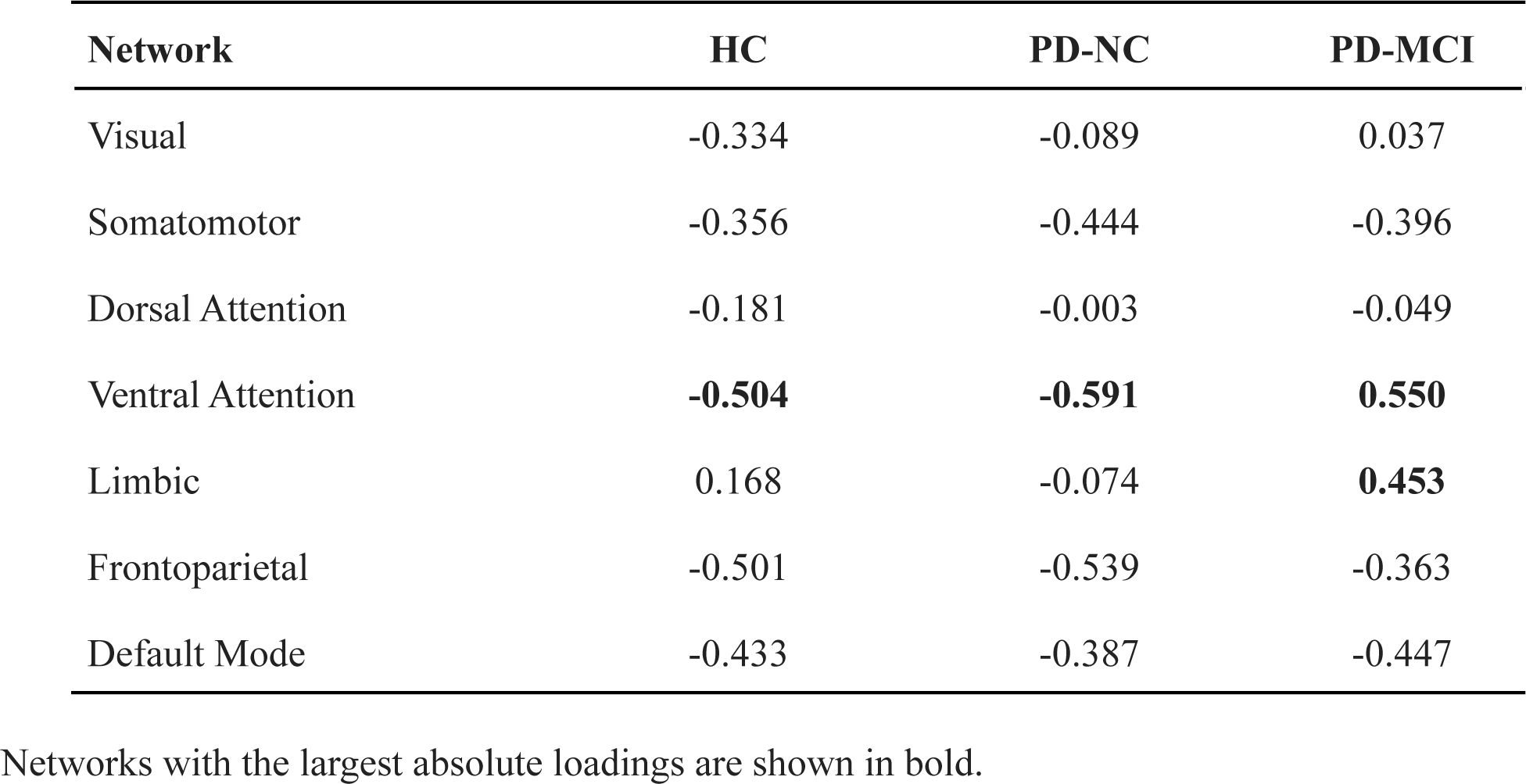
PC1 Loadings of Network-Specific Integration Indices Across Groups.

| Network | HC | PD-NC | PD-MCI |
| --- | --- | --- | --- |
| Visual | -0.334 | -0.089 | 0.037 |
| Somatomotor | -0.356 | -0.444 | -0.396 |
| Dorsal Attention | -0.181 | -0.003 | -0.049 |
| Ventral Attention | **-0.504** | **-0.591** | **0.550** |
| Limbic | 0.168 | -0.074 | **0.453** |
| Frontoparietal | -0.501 | -0.539 | -0.363 |
| Default Mode | -0.433 | -0.387 | -0.447 |

Networks with the largest absolute loadings are shown in bold.

In PD-NC participants, the dominant organizational axis was strongly defined by the VAN (loading=−0.591) and FPN (loading=−0.539), followed by Somatomotor (loading=−0.444) and DMN (loading=−0.387). Visual (loading=−0.089), LIM (loading=−0.074), and DAN (loading=−0.003) contributed minimally.

In contrast, PD-MCI participants demonstrated a marked reorganization of PC1 loadings. The VAN (loading=0.550) and LIM (loading=0.453) networks emerged as the strongest contributors, whereas DMN (loading=−0.447), Somatomotor (loading=−0.396), FPN (loading=−0.363), DAN (loading=−0.049), and Visual (loading=0.037) networks contributed comparatively less.

Notably, the LIM shifted from a weak contributor in PD-NC to one of the strongest contributors in PD-MCI, while VAN remained the dominant network defining the principal organizational axis. This transition indicates a substantial reconfiguration of network architecture associated with cognitive impairment.

### 3.3 Within-Group Variability and Multivariate Dispersion

Across all three groups, variability profiles remained broadly comparable, with no network exhibiting pronounced increases in dispersion in either PD-NC or PD-MCI participants (Figure 4). HC showed the highest variability in LIM (SD=0.104), Visual (SD=0.094), and VAN (SD=0.092) networks, whereas PD-NC participants demonstrated the lowest overall variability across most networks, particularly within DMN (SD=0.054), FPN (SD=0.059), VAN (SD=0.068), and LIM (SD=0.073) systems. PD-MCI participants exhibited intermediate variability levels that remained comparable to those observed in HC.

**Figure 4.**
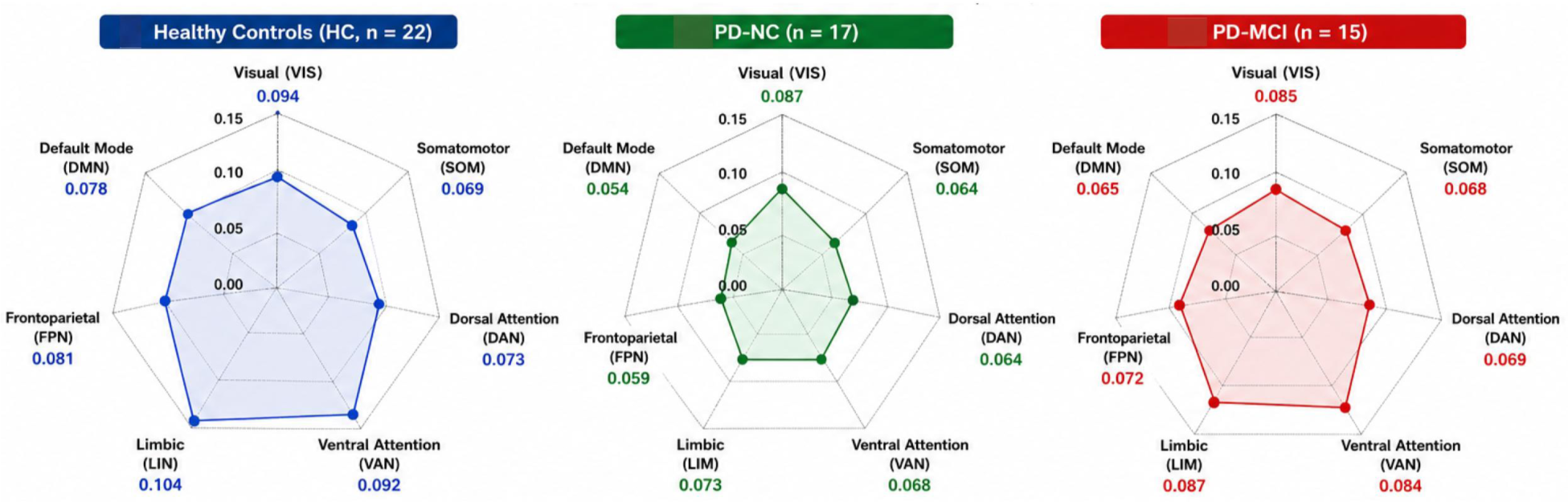
Within-group variability of network-specific integration indices across diagnostic groups. Radar plots show the standard deviation of network-specific integration indices across participants within each group for the seven Yeo networks. HC showed the highest variability in the Limbic, Visual, and Ventral Attention networks, whereas PD-NC showed the lowest overall variability across most networks, particularly within DMN, FPN, VAN, and LIM. PD-MCI showed intermediate variability levels that remained comparable to HC and PD-NC. Global mean standard deviation did not differ significantly across groups, and Dunn–Bonferroni pairwise comparisons were not significant.

Global variability measures did not differ significantly across groups (Kruskal–Wallis: *H*(2)=3.45, *p*=0.178). Pairwise comparisons similarly revealed no significant differences between HC and PD-NC (*p*=0.142), HC and PD-MCI (*p*=0.428), or PD-NC and PD-MCI (*p*=0.356), indicating that the observed alterations in network integration were not accompanied by increased within-group variability.

Mean centroid distances were highly comparable across groups (HC=0.060, PD-NC=0.052, PD-MCI=0.056), and no significant differences were observed (Kruskal–Wallis *p*=0.832). Pairwise comparisons likewise revealed no significant differences between HC and PD-NC (*p*=0.584), HC and PD-MCI (*p*=0.676), or PD-NC and PD-MCI (*p*=0.921).

## 4. Discussion

### 4.1 Conceptual and Methodological Implications of the entropy-based network integration framework

The present study introduced an entropy-based network integration framework for quantifying how canonical resting-state networks are represented across ICA-derived functional modes. Its principal methodological contribution is to characterizes network representational distribution rather than conventional connectivity strength. Empirical findings provided initial evidence that this measure is sensitive to selective group-level differences in large-scale network organization, and convergence across pairwise comparisons, exploratory PCA, variability profiling, and centroid-distance analyses further supports that these differences are not simply explained by increased within-group dispersion. To the best of our knowledge, the proposed metric for ICA is novel and has not been reported in literature.

A key implication is that the framework provides interpretable global and network-specific measures, allowing both overall ICA–network diversity and focal versus distributed representation of canonical systems across ICs to be assessed, respectively. In the present application, group differences in global measures were accompanied by selective network-specific findings, suggesting that the metric can identify organized differences in network representation that may be obscured by global summaries alone.

Conceptually, the framework also extends ICA analysis by converting component–network overlap patterns into quantitative measures of representational entropy and integration. This builds on existing ICA–template mapping practices and enables ICA outputs to be compared statistically across individuals and groups.

### 4.2 Empirical Findings and Biological Interpretation

Multiple analyses reveal a coherent pattern of large-scale network differences across PD cognitive-status groups. Rather than showing uniform alterations across all networks, the findings suggested selective differences involving higher-order cognitive, attentional, and limbic systems. This pattern is generally consistent with prior resting-state functional connectivity studies showing that cognitive impairment in PD is associated with altered connectivity in cognitively relevant large-scale networks, including DMN, FPN, DAN/frontoinsular, salience, and limbic-related systems (Baggio et al., 2015; Amboni et al., 2015; Yeager et al., 2024). The VAN, which overlaps substantially with salience-processing systems, supports the detection of behaviorally relevant stimuli and the reorienting of attention toward salient events (Corbetta & Shulman, 2002; Seeley et al., 2007; Vossel et al., 2014). Higher VAN integration may indicate that attentional network representations are distributed across a broader set of ICA-derived functional modes in PD-MCI, possibly reflecting increased reliance on attentional control mechanisms. In contrast, reduced LIM integration may indicate a more focal or contracted representation in PD-MCI, given the role of limbic systems in emotional regulation, reward processing, motivation, memory formation, and affective decision-making (Mesulam, 1998; Catani et al., 2013; Rolls, 2019), and prior evidence of limbic abnormalities in PD-related cognitive and affective symptoms (Tessitore et al., 2012; Zhan et al., 2018).

The elevated DMN and FPN integration observed in PD-NC participants is notable. The FPN supports adaptive cognitive control and flexible coordination among cognitive systems (Cole et al., 2013; Dixon et al., 2018), whereas the DMN is involved in internally directed cognition, autobiographical memory, self-referential processing, and large-scale information integration (Buckner et al., 2008; Andrews-Hanna et al., 2014). This interpretation is broadly consistent with models of compensatory reorganization in aging and neurodegeneration, and with prior resting-state studies reporting altered executive and DMN involvement in cognitively preserved PD (Olde Dubbelink et al., 2014; Reuter-Lorenz & Park, 2014; Gorges et al., 2015; Baggio et al., 2015; Cabeza et al., 2018). However, because individual-level clinical and cognitive associations were not available, this compensatory interpretation remains preliminary.

The PCA findings provided an important multivariate perspective to these results. While HC and PD-NC participants were primarily characterized by attentional–executive organizational patterns dominated by VAN, FPN, and DMN contributions, PD-MCI participants demonstrated a marked reconfiguration of the dominant organizational axis. Specifically, VAN (loading=0.550) and LIM (loading=0.453) networks emerged as strongest contributors to PC1, indicating convergence across independent analytical approaches. Similar transitions between executive-control systems and salience-attention systems have been proposed as key mechanisms underlying cognitive decline and network reorganization in neurodegenerative disorders (Seeley et al., 2007; Menon, 2011; Jones et al., 2016). Notably, the same networks identified through PCA were also those exhibiting significant pairwise differences, suggesting that cognitive-status differences (PD-NC vs PD-MCI) are associated not merely with changes in individual networks but with reconfiguration of the dominant organizational architecture governing large-scale brain function.

A particularly important observation was the preservation of within-group stability despite significant network-level differences. Radar plot analyses demonstrated comparable variability profiles across groups, while distance-to-centroid analyses revealed no significant differences in multivariate dispersion - suggesting that cognitive-status differences in the present cross-sectional dataset were associated with coherent redistribution of network representations rather than nonspecific increases in noise, variability, or dispersion. Such an interpretation is consistent with contemporary models of neurodegeneration that emphasize systematic network reorganization rather than random network breakdown (Seeley et al., 2007; Stam, 2014; Jones et al., 2016).

### 4.3 Potential Applications for Longitudinal and Clinical Network Tracking

Building on the present findings, the proposed entropy-based integration framework may have broader utility for studying longitudinal changes in large-scale brain organization. It may provide a potential means of tracking whether network representations become more focal, more distributed, or redistributed over time. The proposed framework may therefore offer a complementary approach for characterizing how network representations evolve across developmental stages, clinical trajectories, or therapeutic interventions.

A network representational approach may also be useful for investigating shared and disorder-specific patterns across clinically distinct conditions. Many psychiatric and neurodevelopmental disorders show overlapping symptoms and partially shared abnormalities in default-mode, salience, attention, and executive-control systems. By quantifying the representational distribution of networks rather than focusing only on pairwise connectivity strength, entropy-derived integration indices may help characterize whether different conditions involve similar or distinct patterns of network reorganization. Such applications may contribute to a more network-oriented understanding of clinical heterogeneity and transdiagnostic mechanisms.

These possibilities remain speculative and require validation in larger longitudinal datasets with behavioral and clinical measures.

### 4.4 Limitations and Future Directions

Several limitations should be considered when interpreting the present findings. First, although the framework was applied to an open dataset included HC, PD-NC, and PD-MCI participants, the sample size within each group remained modest. Larger cohorts are needed to establish the robustness and generalizability of the observed entropy-derived integration profiles across populations, disease stages and acquisition settings.

Second, the framework was evaluated using rs-fMRI ICA-derived network decompositions mapped onto the Yeo seven-network atlas. Although this provides a standardized representation of large-scale brain organization, future studies should examine the stability of the findings across alternative parcellation schemes, higher-resolution network definitions, and different ICA model orders.

Third, the used dataset was cross-sectional and lacked individual-level cognitive performance scores, disease-severity measures, disease-duration, and medication-related variables. Therefore, the findings should be interpreted as evidence that the proposed metric is sensitive to group-level cognitive-status differences, rather than as evidence that it tracks cognitive decline, disease severity, or clinical progression at the individual-subject level. Longitudinal studies with detailed neuropsychological and clinical measures will be necessary to evaluate these possibilities.

Fourth, direct comparisons with established measures, including graph-theoretical metrics, functional connectivity strength, dynamic functional connectivity, spatial similarity or overlap, and signal-based entropy measures, would help clarify the unique and incremental information captured by the proposed framework.

Fifth, formal test-retest reliability could not be assessed because the dataset included only a single rs-fMRI run per participant. Split-half reliability was also not considered appropriate because dividing the available resting-state acquisition into shorter segments would likely provide insufficient temporal information for stable independent component estimation, particularly when using automatic dimensionality estimation. Future validation studies using repeated resting-state acquisitions or longer scan durations are therefore needed to estimate the reliability of the proposed indices.

## 5. Conclusion

The current study provides evidence that entropy-derived network representational integration offers a novel perspective on large-scale brain organization and may complement existing approaches for characterizing functional network architecture in health and disease.

Findings from our proposed metric suggest a cross-sectional pattern in which cognitively preserved PD was associated with enhanced DMN–FPN integration, whereas PD-MCI was characterized by selective VAN–LIM reconfiguration. These cognitive-status differences may involve a redistribution of network representational architecture in which attentional systems become increasingly prominent while limbic contributions are altered. The convergence of results across multiple independent analytical approaches supports the robustness of this interpretation.

Because the used dataset did not include repeated resting-state scans or individual-level clinical severity measures, the present analysis should be interpreted as a preliminary validation of group-level sensitivity rather than a complete validation of reliability, clinical prediction, or individual-level disease tracking.

## Statements and Declarations

### Funding and/or Competing Interests

No funding was received for this study. The authors have no relevant financial or non-financial interests to disclose.

### Ethics approval

This study used openly available, de-identified resting-state fMRI data obtained from OpenNeuro. No new human participants were recruited, and no new data were collected for the present analysis. Because the present work involved secondary analysis of publicly available de-identified data, separate institutional ethics approval was not required.

### Data Availability Statement

The resting-state fMRI dataset analyzed in this study is publicly available through OpenNeuro under dataset accession number ds005892. The present study used this open dataset for secondary analysis. Derived data, analysis scripts, and entropy-derived integration indices generated during the present study will be made available by the corresponding author upon reasonable request.

### Author Contributions

CRediT Taxonomy. Conceptualization and Design: Poulami Kar; Literature Search: Poulami Kar, Dipayan Roy; Writing - Formulae Development: Poulami Kar; Writing - data acquisition, analysis and interpretation: Poulami Kar, Dipayan Roy; Writing-original draft preparation: Poulami Kar, Dipayan Roy; Writing - review and editing: Bhoomika R. Kar, Dipayan Roy; Writing - illustrations: Poulami Kar; Supervision: Bhoomika R. Kar All authors read and approved the final version of the manuscript.

### Clinical trial number

Not applicable. *Consent to Participate:* Not applicable. *Consent to Publish:* Not applicable.

## References

Amboni, M., Tessitore, A., Esposito, F., Santangelo, G., Picillo, M., Vitale, C., Giordano, A., Erro, R., de Micco, R., Corbo, D., & Barone, P. (2015). Resting-state functional connectivity associated with mild cognitive impairment in Parkinson’s disease. Journal of Neurology, 262(2), 425–434. 10.1007/s00415-014-7591-5

Andrews-Hanna, J. R., Smallwood, J., & Spreng, R. N. (2014). The default network and self-generated thought: Component processes, dynamic control, and clinical relevance. Annals of the New York Academy of Sciences, 1316(1), 29–52. 10.1111/nyas.12360

Baggio, H. C., Segura, B., Sala-Llonch, R., Marti, M. J., Valldeoriola, F., Compta, Y., Tolosa, E., & Junqué, C. (2015). Cognitive impairment and resting-state network connectivity in Parkinson’s disease. Human Brain Mapping, 36(1), 199–212. 10.1002/hbm.22622

Bassett, D. S., & Sporns, O. (2017). Network neuroscience. Nature Neuroscience, 20(3), 353–364. 10.1038/nn.4502

Buckner, R. L., Andrews-Hanna, J. R., & Schacter, D. L. (2008). The brain’s default network: Anatomy, function, and relevance to disease. Annals of the New York Academy of Sciences, 1124(1), 1–38. 10.1196/annals.1440.011

Bullmore, E., & Sporns, O. (2009). Complex brain networks: Graph theoretical analysis of structural and functional systems. Nature Reviews Neuroscience, 10(3), 186–198. 10.1038/nrn2575

Cabeza, R., Albert, M., Belleville, S., Craik, F. I. M., Duarte, A., Grady, C. L., Lindenberger, U., Nyberg, L., Park, D. C., Reuter-Lorenz, P. A., Rugg, M. D., Steffener, J., & Rajah, M. N. (2018). Maintenance, reserve and compensation: The cognitive neuroscience of healthy ageing. Nature Reviews Neuroscience, 19(11), 701–710. 10.1038/s41583-018-0068-2

Catani, M., Dell’Acqua, F., & De Schotten, M. T. (2013). A revised limbic system model for memory, emotion and behaviour. Neuroscience & Biobehavioral Reviews, 37(8), 1724–1737. 10.1016/j.neubiorev.2013.07.001

Cole, M. W., Reynolds, J. R., Power, J. D., Repovs, G., Anticevic, A., & Braver, T. S. (2013). Multi-task connectivity reveals flexible hubs for adaptive task control. Nature Neuroscience, 16(9), 1348–1355. 10.1038/nn.3470

Corbetta, M., & Shulman, G. L. (2002). Control of goal-directed and stimulus-driven attention in the brain. Nature Reviews Neuroscience, 3(3), 201–215. 10.1038/nrn755

Dixon, M. L., De La Vega, A., Mills, C., Andrews-Hanna, J., Spreng, R. N., Cole, M. W., & Christoff, K. (2018). Heterogeneity within the frontoparietal control network and its relationship to the default and dorsal attention networks. Proceedings of the National Academy of Sciences, 115(7), E1598–E1607. 10.1073/pnas.1715766115

Garrett, D. D., Kovacevic, N., McIntosh, A. R., & Grady, C. L. (2011). The importance of being variable. The Journal of neuroscience: the official journal of the Society for Neuroscience, 31(12), 4496–4503. 10.1523/JNEUROSCI.5641-10.2011

Gorges, M., Müller, H. P., Lulé, D., Del Tredici, K., Ludolph, A. C., Pinkhardt, E. H., & Kassubek, J. (2015). To rise and to fall: Functional connectivity in cognitively normal and cognitively impaired patients with Parkinson’s disease. Neurobiology of Aging, 36(4), 1727–1735. 10.1016/j.neurobiolaging.2014.12.026

Jones, D. T., Knopman, D. S., Gunter, J. L., Graff-Radford, J., Vemuri, P., Boeve, B. F., Petersen, R. C., Weiner, M. W., Jack, C. R., & Alzheimer’s Disease Neuroimaging Initiative. (2016). Cascading network failure across the Alzheimer’s disease spectrum. Brain, 139(2), 547–562. 10.1093/brain/awv338

Kemp, A. S., Eubank, J., Younus, Y., Galvin, J. E., Prior, F. W., & Larson-Prior, L. J. (2025). Resting state MRI data from healthy controls (HC), Parkinson’s disease with normal cognition (PD-NC), and Parkinson’s disease with mild cognitive impairment (PD-MCI) cohorts [Data set]. OpenNeuro. 10.18112/openneuro.ds005892.v1.0.0

Lesiuk, T., Bugos, J. A., & Murakami, B. (2018). A Rationale for Music Training to Enhance Executive Functions in Parkinson’s Disease: An Overview of the Problem. Healthcare (Basel, Switzerland), 6(2), 35. 10.3390/healthcare6020035

McIntosh, A. R., Kovacevic, N., Lippe, S., Garrett, D., Grady, C., & Jirsa, V. (2010). The development of a noisy brain. Archives italiennes de biologie, 148(3), 323–337.

Menon, V. (2011). Large-scale brain networks and psychopathology: A unifying triple network model. Trends in Cognitive Sciences, 15(10), 483–506. 10.1016/j.tics.2011.08.003

Mesulam, M. M. (1998). From sensation to cognition. Brain, 121(6), 1013–1052. 10.1093/brain/121.6.1013

Niu, Y., Wang, B., Zhou, M., Xue, J., Shapour, H., Cao, R., Cui, X., Wu, J., & Xiang, J. (2018). Dynamic Complexity of Spontaneous BOLD Activity in Alzheimer’s Disease and Mild Cognitive Impairment Using Multiscale Entropy Analysis. Frontiers in neuroscience, 12, 677. 10.3389/fnins.2018.00677

Nomi, J. S., Farrant, K., Damaraju, E., Rachakonda, S., Calhoun, V. D., & Uddin, L. Q. (2016). Dynamic functional network connectivity reveals unique and overlapping profiles of insula subdivisions. Human brain mapping, 37(5), 1770–1787. 10.1002/hbm.23135

Olde Dubbelink, K. T. E., Hillebrand, A., Stoffers, D., Deijen, J. B., Twisk, J. W. R., Stam, C. J., & Berendse, H. W. (2014). Disrupted brain network topology in Parkinson’s disease: A longitudinal magnetoencephalography study. Brain, 137(1), 197–207. 10.1093/brain/awt316

Park, C. H., Chang, W. H., Ohn, S. H., Kim, S. T., Bang, O. Y., Pascual-Leone, A., & Kim, Y. H. (2011). Longitudinal changes of resting-state functional connectivity during motor recovery after stroke. Stroke, 42(5), 1357–1362. 10.1161/STROKEAHA.110.596155

Reuter-Lorenz, P. A., & Park, D. C. (2014). How does it STAC up? Revisiting the scaffolding theory of aging and cognition. Neuropsychology Review, 24(3), 355–370. 10.1007/s11065-014-9270-9

Rolls, E. T. (2019). The cingulate cortex and limbic systems for emotion, action, and memory. Brain Structure and Function, 224(9), 3001–3018. 10.1007/s00429-019-01945-2

Shannon CE. A mathematical theory of communication. Bell Syst Tech J. 1948;27:379–423.

Seeley, W. W., Menon, V., Schatzberg, A. F., Keller, J., Glover, G. H., Kenna, H., Reiss, A. L., & Greicius, M. D. (2007). Dissociable intrinsic connectivity networks for salience processing and executive control. Journal of Neuroscience, 27(9), 2349–2356. 10.1523/JNEUROSCI.5587-06.2007

Stam, C. J. (2014). Modern network science of neurological disorders. Nature Reviews Neuroscience, 15(10), 683–695. 10.1038/nrn3801

Tessitore, A., Esposito, F., Vitale, C., Santangelo, G., Amboni, M., Russo, A., Corbo, D., Cirillo, G., & Barone, P. (2012). Default-mode network connectivity in cognitively unimpaired patients with Parkinson disease. Neurology, 79(23), 2226–2232. 10.1212/WNL.0b013e31827689d6

Vossel, S., Geng, J. J., & Fink, G. R. (2014). Dorsal and ventral attention systems. Neuroscientist, 20(2), 150–159. 10.1177/1073858413494269

Wang, Z., Li, Y., Childress, A. R., & Detre, J. A. (2014). Brain entropy mapping using fMRI. PloS one, 9(3), e89948. 10.1371/journal.pone.0089948

Yeager, B. E., Twedt, H. P., Bruss, J., Schultz, J., & Narayanan, N. S. (2024). Cortical and subcortical functional connectivity and cognitive impairment in Parkinson’s disease. NeuroImage. Clinical, 42, 103610. 10.1016/j.nicl.2024.103610

Yeo, B. T., Krienen, F. M., Sepulcre, J., Sabuncu, M. R., Lashkari, D., Hollinshead, M., Roffman, J. L., Smoller, J. W., Zöllei, L., Polimeni, J. R., Fischl, B., Liu, H., & Buckner, R. L. (2011). The organization of the human cerebral cortex estimated by intrinsic functional connectivity. Journal of neurophysiology, 106(3), 1125–1165. 10.1152/jn.00338.2011

Zhan, Z. W., Lin, L. Z., Yu, E. H., Xin, J. W., Lin, H. L., & Li, Y. C. (2018). Abnormal resting-state functional connectivity in Parkinson’s disease with mild cognitive impairment. CNS Neuroscience & Therapeutics, 24(3), 188–194. 10.1111/cns.12769

